# Derivation and simulation of a computational model of active cell populations: How overlap avoidance, deformability, cell-cell junctions and cytoskeletal forces affect alignment

**DOI:** 10.1101/2024.02.02.578535

**Authors:** Vivienne Leech, Fiona N Kenny, Stefania Marcotti, Tanya J Shaw, Brian M Stramer, Angelika Manhart

## Abstract

Collective alignment of cell populations is a commonly observed phenomena in biology. An important example are aligning fibroblasts in healthy or scar tissue. In this work we derive and simulate a mechanistic agent-based model of the collective behaviour of actively moving and interacting cells, with a focus on understanding collective alignment. The derivation strategy is based on energy minimisation. The model ingredients are motivated by data on the behaviour of different populations of aligning fibroblasts and include: Self-propulsion, overlap avoidance, deformability, cell-cell junctions and cytoskeletal forces. We find that there is an optimal ratio of self-propulsion speed and overlap avoidance that maximises collective alignment. Further we find that deformability aids alignment, and that cell-cell junctions by themselves hinder alignment. However, if cytoskeletal forces are transmitted via cell-cell junctions we observe strong collective alignment over large spatial scales.

## 1 Introduction

### The challenge of active particles

The ability of particles to align with their neighbours is observed in many contexts and in many scales in biology. Famous examples include flocking of birds, schools of fish or even motion of large groups of people [1, 2, 3]. Here we focus on alignment on a cellular scale. Cell alignment has been observed in bacterial swarms [4], e.g. in myxobacteria [5], as well as in amoeboid cells [6], or fibroblasts [7]. Alignment of different types of particles has also been observed and modelled in physics [8, 9, 10], however alignment in cells is more challenging to understand, since cells can exhibit much more complicated behaviour than passive matter, see e.g. reviews [11, 12]. In particular, cells can actively self-propel, can change their shape, can interact biochemically via signalling and mechanically via physical junctions with their neighbours and with the extracellular matrix, with several levels of potential feedback involved.

### Causes of alignment

Larger, more complex organisms, such as birds or fish, are typically able to perceive their neighbours using senses such as sight and can adjust their own direction and speed correspondingly. Some interesting work for causes of alignment for such species include [13, 14]. Cells, the main focus of this work, can perceive their surroundings in several different ways. The main ones include chemotaxis, durotaxis, signalling or mechanics. Several experimental works [15, 16] have shown that e.g. the extracellular matrix (ECM) can transmit forces, aiding cytoskeletal alignment. This has been modelled e.g. in [17, 18]. Further, the shape of the environment itself can influence alignment by introducing constraints to cell orientation on the boundary [7]. Finally, contact-based alignment can be caused by particles avoiding overlap. For movement in e.g. a fluid this would be a hard constraint, while for crawling cells “overlap” could imply cells moving on top of each other.

### Basic types of alignment models

Dynamics and patterns emerging from the interactions of many individuals are difficult to intuit from microscopic interaction rules, hence mathematical modelling and simulations are a powerful tool to shed light on the involved mechanisms. One common model type is continuum models, where the system is described in terms of continuous space and time dependent macroscopic quantities like cell density, mean direction, etc. The book [19] offers an excellent overview of models and biomedical applications. Continuum models have the advantage of ease of analysis and have been used extensively to investigate alignment and pattern formation in cell populations, for some examples see [20, 21, 22, 23, 24, 25, 26]. Agent-based models, on the other hand, where cells are discrete objects (“agents”) and each cell is equipped with its own set of equations, are particularly suitable for mechanistic hypothesis testing, since biological assumptions can be translated in a relatively straight-forward way (see e.g. reviews [27, 28]). Amongst the classical agent-based models for flocking ([29, 30]), the most famous agent-based alignment model is probably the Vicsek model [31]. This model assumes self-propelling particles align their orientation with their neighbours and produces large scale alignment whenever the alignment force is large compared to the orientational noise. While the Vicsek model and its variants have been applied to many biological problems [32, 33], it isn’t suitable to test mechanisms of alignment, since alignment is already a model ingredient.

### Shape deformations

For individual cells or small groups of cells, there exist several modelling frameworks capable of describing cell shapes in a flexible and biologically well-motivated manner. Prominent approaches include 1. Phase-field models [34, 35, 36, 37, 38], where the cell in- and outsides are characterised by a continuous, but steep phase-field variable, 2. Models using the immersed boundary method [39, 40, 41], where the cell boundary is an explicit curve or surface interacting with the surrounding fluid, 3. Cellular potts models [42, 43, 44], where each cell is a collection of pixels, whose dynamics follow an energy minimization, or 4. Vertex models [45, 46], which describe sheets of cells via cell-cell boundaries. While phase-field, immersed boundary and cellular potts models allow for the description of a large class of cell shapes, they are also computationally costly and hence less suitable to investigate large numbers of cells. Vertex models, on the other hand, are mostly used for simulating tissue dynamics and are less suitable for describing individual cell movement.

### This work and paper overview

In this work, we will mathematically model alignment of collectives of cells moving in two space dimensions (2D). As the main cause of alignment, we assume cells want to avoid overlap in a tuneable (i.e. non-perfect) way (Sec. 2). Motivated by biologically observed, distinct behaviour of different populations of fibroblasts [47], we will assess the influence of the following characteristics of active matter on alignment: Self-propulsion (Sec. 2), deformability (Sec. 3), cell-cell junctions (Sec. 4) and cytoskeletal forces (Sec. 4).

## 2 The base model: Self-propulsion and overlap avoidance

### 2.1 Biological background & model ingredients

#### Experimental motivation

While alignment processes in active particles are relevant in many contexts, we will focus on the particular example of fibroblasts. Fibroblasts are cells in the connective tissue in animals and are responsible for making and remodelling the ECM. Fibroblast and ECM alignment is observed during various scarring pathologies. In [47] we have investigated the difference in alignment behaviour of fibroblasts in healthy tissue (normal dermal fibroblasts, NDFs) as compared to dermofibroblasts in certain scar tissue (keloid derived fibroblasts, KDFs). KDFs were found to show stronger alignment over larger length scales, Fig. 1A,B. Further we found that KDFs show less tendency to crawl on top of each other and form aligned supracellular actin bundles via cell-cell junctions, spanning multiple cells. Using mathematical modelling, we found in [47] that the increase in overlap avoidance can explain the stronger alignment. In this work we will use the differences found in NDFs and KDFs to motivate further model extensions. However, the model ingredients are applicable to other cells and situations and hence the findings are relevant beyond fibroblasts.

**Figure 1:**
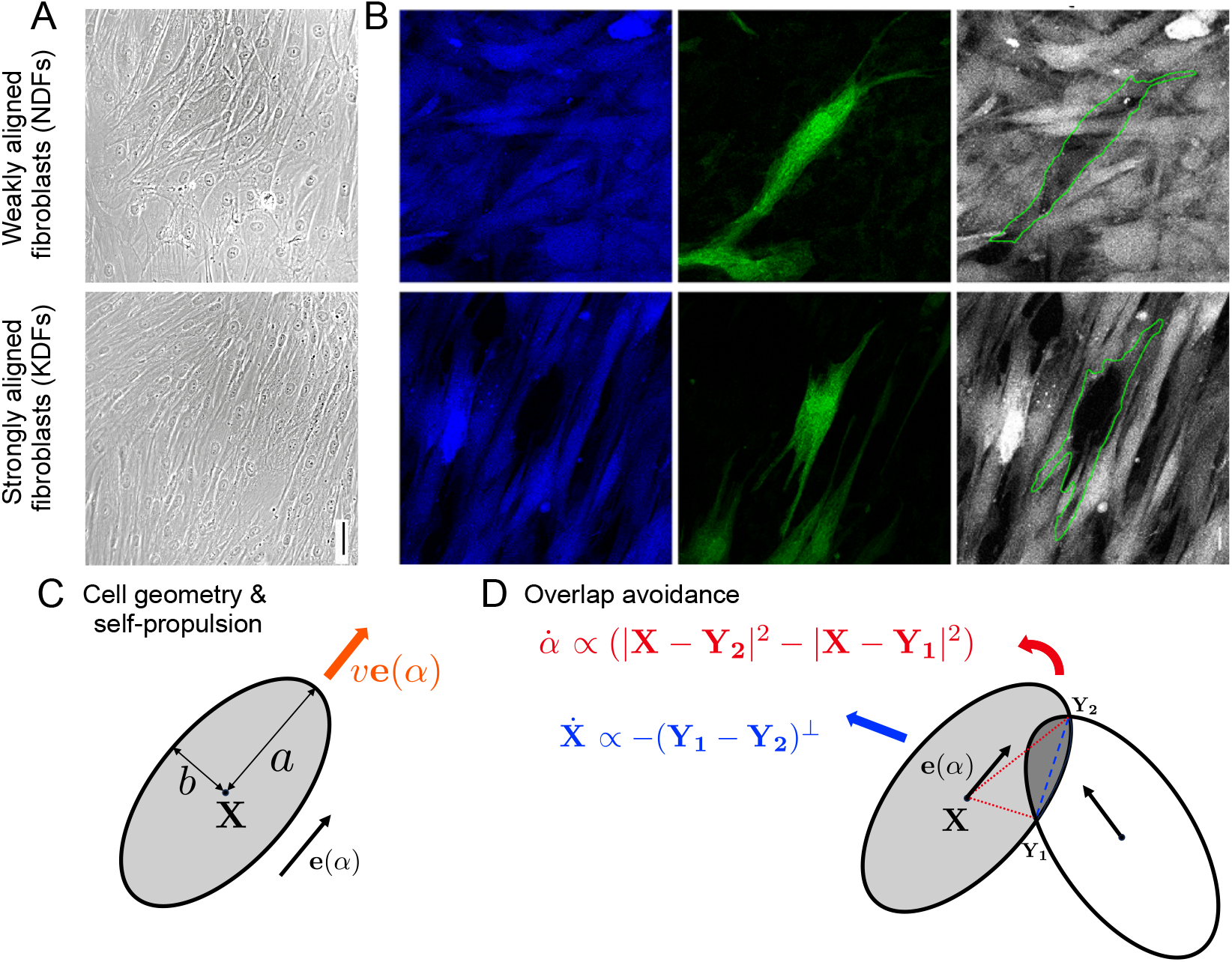
A,B: Experimental figures for weakly aligned normal dermal fibroblasts (NDFs), top row, and strongly aligned keloid derived fibroblasts (KDFs), bottom row. A: Phase microscopy pictures, scale bar 50 *μ*m. B: Mosaically labelled cells with two different probes (left: CellTrace Violet, middle: CellTrace Green, right: overlay), scale bar 20 *μ*m. C,D: Model schematics. C: Elliptic cell geometry of a single cell with centre **X**, dimensions *a* and *b*, orientation *α* and self-propulsion speed *v*. D: Effect of overlap avoidance upon one cell overlapping with another cell.

#### Model ingredients

We build a mechanistic, agent-based model to describe the motion of individual cells interacting with neighbouring cells in 2D. While cell shapes can be very complicated and include protrusions, ruffles, etc., we approximate cells as ellipses. This captures the fact that actively moving cells like fibroblasts are typically spindle-shaped and leads to an easier model formulation while allowing for cell deformations (see Sec. 3). In this work we do not model cell divisions, however, they could be included in a straight-forward manner.

The key model ingredients are

- *Environmental friction:* As usual in cell biology, we assume a friction-dominated regime. As a consequence, velocities (not accelerations) are proportional to forces. The strength of the friction with the substrate is given by *η*, which effectively sets a time scale.
- *Self-propulsion:* In the absence of interactions, cells move with fixed speed *v* in the direction of their orientation. Orientational noise could be included in a straight forward manner, but is omitted in this work for the sake of simplicity.
- *Overlap avoidance:* When placed on a 2D substrate, many cells types will tend to avoid moving on top of other cells. The term *contact inhibition of locomotion* is sometimes used in this context. However, contact inhibition of locomotion more commonly refers to an active change of direction upon contact, as opposed to a more passive reaction, which is what we model here. Another commonly used term in this context is *repulsion*, however, in this work will use the term *overlap avoidance* to emphasise that the effect is short-ranged and driven by cell overlap. Note that “overlap” in 2D can be interpreted either as being positioned partly on top of each other, or allowing for some cell softness. We allow overlap avoidance to be tuneable, its strength given by parameter *σ*. If *σ* = 0, cells have no overlap avoidance, and for *σ → ∞*, cells would behave as solid objects that never overlap/move on top of each other. To avoid overlap cells can
  - move to change their location,
  - turn to change their orientation, or
  - change their shape (*→* Sec. 3).
  All these effects will be a consequence of the minimisation of a common energy term.
- *Cell-cell junctions:* In Sec. 4 we model and investigate the effect of cell-cell junctions, where cells are elastically tethered to each other, which can affect their orientation and position.
- *Actin forces:* Also in Sec. 4 we describe the presumed effect of supracellular actin cables that lead to cytoskeletal forces affecting cell orientation.

### 2.2 Model derivation

We will show the derivation excluding cell deformability and cell-cell junctions. These will be considered in Sec. 3 and Sec. 4. More derivation details can be found in S1 Appendix, Sec. 1 and a summary of model parameter names and meaning can be found in S1 Appendix Tab. 1. We consider *N* cells within the fixed domain Ω *∈* ℝ^2^, each with position **X**_*i*_ = (*X*_*i*_, *Y*_*i*_) *∈* ℝ^2^, *i* = 1, …, *N* and orientation *α*_*i*_ *∈* [0, 2*π*), *i* = 1, …, *N*. Each cell is described by an ellipse with semi-major axis *a* and semi-minor axis *b* as shown in Fig. 1C. The cell’s area is given by *A* = *abπ*. In the absence of other cells, each cell self-propels with constant velocity *ν* in direction **e**(*α*_*i*_) = (cos(*α*_*i*_), sin(*α*_*i*_))^*T*^, where superscript *T* denotes the transpose.

We derive the governing equations using energy minimisation. Focusing on one cell positioned at **X**(*t*) with orientation *α*(*t*) at time *t*, we parameterise the points inside the cell by

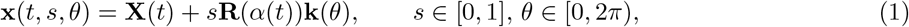

where the rotation matrix **R**(*α*) and the shape vector **k**(*θ*) are defined by

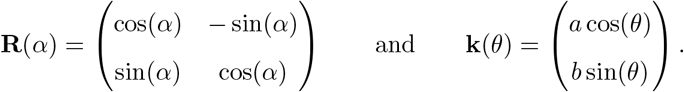

We assume that at every time step Δ*t*, the system minimizes a total energy *E*_tot_, which, for the base model, is the sum of contributions from friction *E*_friction_, from overlap avoidance *E*_overlap_ and from self-propulsion *E*_prop_. All terms are formulated as integrals over the whole (elliptic) cell area using the parametrization given in (1), leading to the area elements *abs. E*_friction_ models friction with the environment by comparing how much points have moved between time *t* and time *t −* Δ*t*:

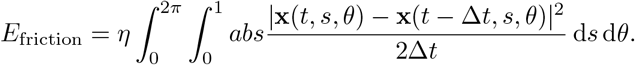

The overlap avoidance term *E*_overlap_ is modelled by an energy potential *V* which includes interactions with all other cells. The choice of *V* will be discussed below.

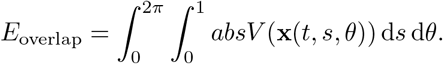

Finally, we want to model self-propulsion. One way to do this is to include it in the energy formulation by prescribing a force **F** acting on the cell. In the course of the derivation it is chosen to be **F** = *νη***e**(*α*), i.e. acting in the direction of the orientation and proportional to the experienced friction *η*. This choice of **F** leads to a self-propulsion speed that is independent of friction.

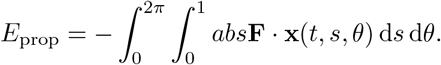

The total energy is then given by

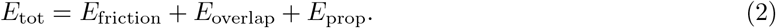

We obtain governing equations by minimising this energy in each time step. In other words, in each time step the cell can change its characteristics (position, orientation, shape) to decrease the energy. Calculation details can be found in S1 Appendix Sec. 1. The main derivation steps for the base model are: 1. Differentiation with respect to **X** and *α* respectively (treating all other variables in the energy potential as constants). 2. Setting the derivative to zero and taking the limit Δ*t →* 0. 3. Evaluation of the integrals. We then obtain the following differential equations for the motion of one cell

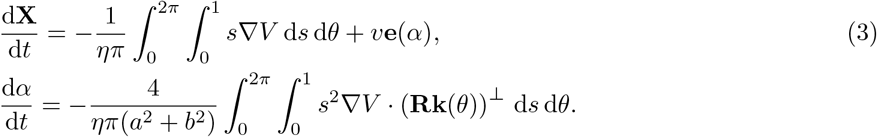

The superscript ⊥ describes the left-turned normal vector. The two equations show how the position and orientations are influenced by the force and torque associated with *V* respectively. Note that we are working in a friction-dominated regime, which is why velocity and angular velocity (as opposed to acceleration and angular acceleration) are proportional to force and torque.

#### Choice of overlap potential *V*

The potential *V* describes the influence of overlap, where *V >* 0 describes overlap avoidance and *V <* 0 overlap preference. Many choices of *V* are possible: e.g. since cells might be thicker closer to the cell center, overlap closer to the cell center could be punished more than further away. However, complicated shapes of *V* are computationally harder to evaluate, especially in the context of collective dynamics. We therefore choose *V* to be constant with value *σ* in regions of overlap and zero elsewhere: For two overlapping ellipses with domains *𝒜* and *ℬ* we define 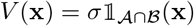, where 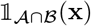 is the indicator function which equals 1 if **x** *∈ 𝒜 ∩ ℬ* and 0 otherwise. The strength of this potential is *σ ∈* ℝ. If *σ >* 0, the cells experience repulsion in response to overlap, and if *σ <* 0, the cells experience attraction. In this work *σ >* 0. Two cells only experience overlap avoidance upon overlapping with each other, hence we define *𝒩*_*i*_ as the set of indices of cells that overlap with the *i*-th cell.

#### Final base model

We non-dimensionalise the model using as reference time 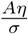, as reference length 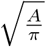 and define *r* = *a/b* as the cell’s aspect ratio. The above choice of *V* allows to evaluate the integrals in (3) explicitly (for calculation details see S1 Appendix, Sec. 1). The resulting equations can be formulated such that they depend only on the points of overlap between cells *i* and *j*, denoted by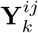, where up to *k* = 4 points of overlap are possible. In the following *K*_*ij*_ = 1 or *K*_*ij*_ = 2 denotes the number of overlap point pairs between cell *i* and cell *j* (having one or three points of overlap can be reduced to having zero or two points of overlap). The overlap points are ordered such that they traverse the boundary of the cell in an anti-clockwise direction. 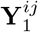 and 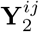 (and 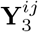 and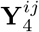) are chosen in such a way that the boundary segment of cell *I* between these pairs of points is contained in the domain of cell *j*, see Fig. 1D.

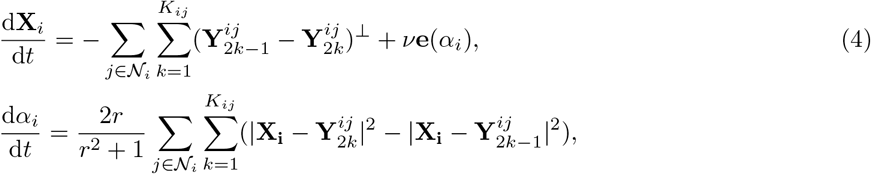

where the non-dimensional quantity *ν* is given by 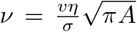 and can be interpreted as comparing the strength of repulsion in the presence of friction to the self-propulsion speed (see more interpretation in the results section below). These two governing equations are supplemented with initial conditions and boundary conditions. Throughout this work the domain is a square box with side length *L* and cells are initially placed randomly inside the box with a random orientation. Further, we use periodic boundary conditions.

#### Interpretation for two cells

To understand the equations better, we consider a situation where there is only interaction between one cell with center **X** and orientation *α*, and one other cell. If there is only one pair of overlap points, **Y**_1_ and **Y**_2_, then (4) reduces to

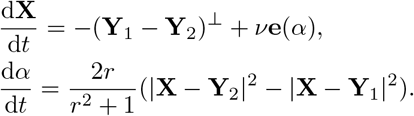

We see that the cell’s center is being pushed in the direction normal to the vector connecting the points of overlap. Further, the change in orientation depends on the difference in lengths of the segments connecting the cell center with the intersection points, turning the cell in the direction from the shorter to the longer one, see Fig. 1D.

### 2.3 Results 1: The base model

Computational details can be found in S1 Appendix, Sec. 2.

#### Alignment increases over time and for larger aspect ratios

We start by demonstrating basic model behaviour. In Fig. 2A,B we see the typical behaviour of the base model with overlap avoidance. Over time cell overlap decreases and alignment increases until a dynamic equilibrium is reached. At this point cells still move, but the alignment parameter stays relatively constant. Further, we observe that cells tend to be aligned with their direct neighbours, but this alignment is local and doesn’t typically go beyond one or two cell lengths. This is related to packing problems, where one studies how and how densely objects of a certain shape can be placed in space without overlapping. Such problems are highly non-trivial, but well-studied for non-moving, completely solid particles, and for symmetric and elongated particles in 2D and 3D (see e.g. [8, 9, 10]). For applications to active matter and in biology see e.g. [17, 48, 49, 50]. Comparison with these works shows that, irrespective of the model details, overlap avoidance leads to some amount of alignment. Next we inspect how the aspect ratio impacts alignment. We find that increasing the aspect ratio *r* of the cells leads to increased alignment. This can be seen in Fig. 2C,D. Investigating this further, we see in Fig. 2C that increasing the aspect ratio of the cells also leads to a higher packing fraction, meaning that there is less cell overlap in the population. Interestingly, for non-moving, solid ellipses [10] found a similar dependence of the alignment parameter on the aspect ratio, albeit with a peak near *r ≈* 1.3.

**Figure 2:**
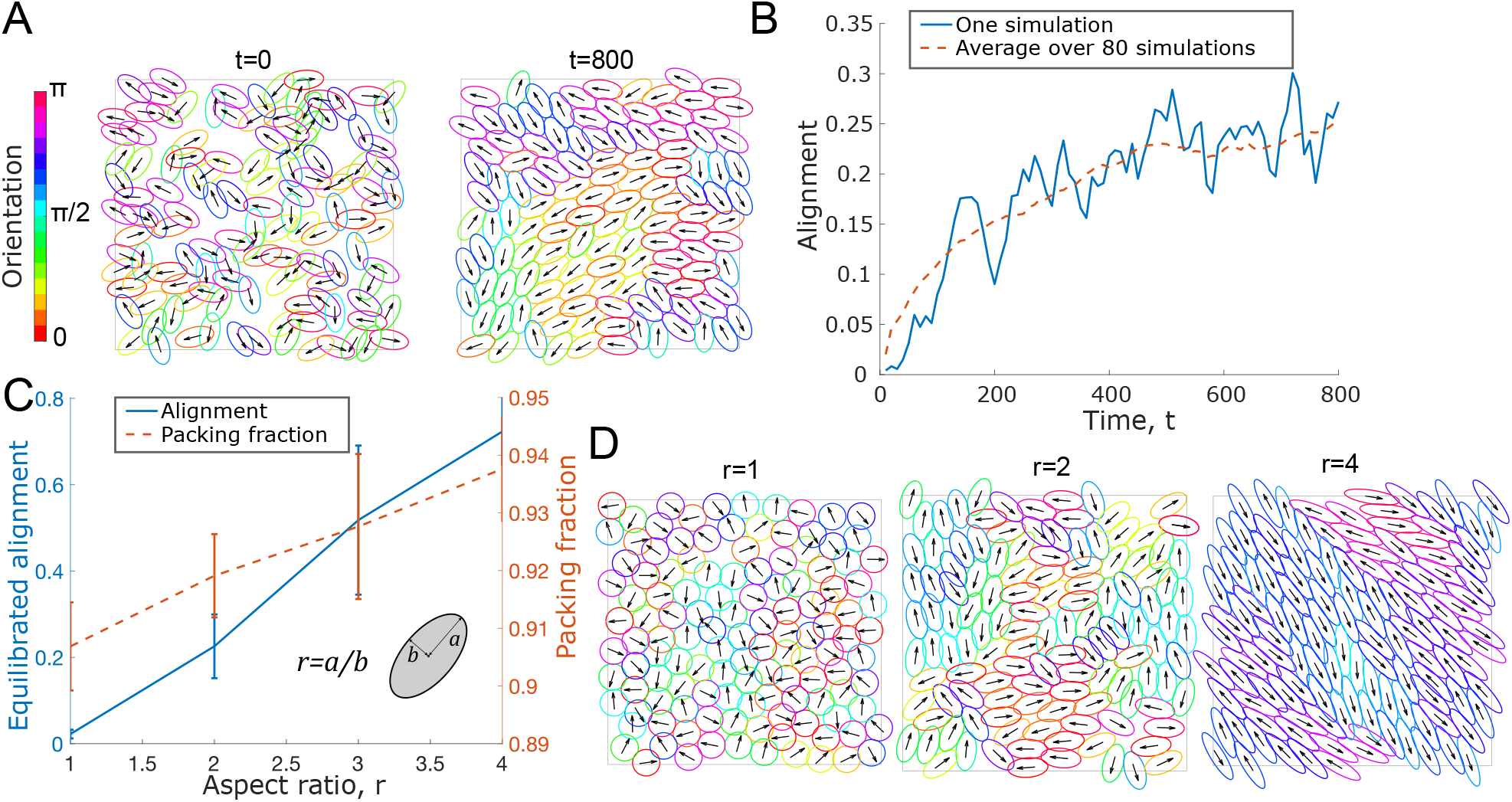
A: Simulation snapshots at time *t* = 0 and *t* = 800 showing cells for an example simulation, color indicates nematic orientation, arrows indicate orientation. See also S2 Video. B: Alignment parameter over time as defined in S1 Appendix, Eq. (1) shown for one individual simulation (blue, solid) and averaged over 80 simulations (orange, dashed). Parameters for A,B: *ν* = 0.2, *N* = 125, *L* = 20, *r* = 2. C: Alignment parameter and packing fraction (see S1 Appendix, Sec. 2), measured at the dynamic equilibrium, plotted against the cell aspect ratio *r*, averaged over 60 simulations, error bars represent standard deviation. D: Simulation snapshot at the (final) time points for three different aspect ratios. Parameters for C,D: *ν* = 0.5, *N* = 125, *L* = 20.

#### Optimal ratio of cell speed to overlap avoidance for alignment

The non-dimensional parameter *ν* is proportional to the self-propulsion speed divided by the strength of overlap avoidance, 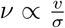, and can be interpreted as the ratio between two time scales *t*_1_*/t*_2_, where 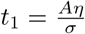 is the time scale of the movement caused by overlap avoidance acting against friction. The time scale 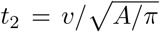 is the time it takes a self-propelling cell of speed *v* to move one reference length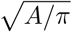. Varying *ν* and measuring the resulting alignment parameter at the dynamic equilibrium, we find a non-monotone dependence on *ν*, with a maximal alignment at *ν* = 0.2, see Fig. 3A,E. We hypothesised that if there is too little self-propulsion, overlap avoidance pushes cells into a little or no overlap configuration, after which cells do not move much and the alignment doesn’t increase further. In that situation, pairs of cells might therefore interact with each other for a long duration, but each cell doesn’t interact with a large number of cells. Self-propulsion, on the other hand, might lead to a re-shuffling of cell contacts, leading to shorter-lived, but more numerous interactions. To test this, we quantified the number of cell contacts over 100 time points for a duration of t=10, and the typical interaction duration (see S1 Appendix, Sec. 2 for details on the quantification). Indeed, we found that the number of interaction partners increases with *ν* (Fig. 3B) and that the interaction duration decreases with *ν* (Fig. 3C). This leads us to suggest the following explanation for the optimal value *ν* found in Fig. 3A: Effective alignment requires cells: 1. To be in contact with their neighbours over a sufficiently long duration (for overlap avoidance to take effect and cause cells to re-orient locally, time scale given by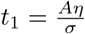) and 2. To be in contact with sufficiently numerous different cells (time scale given by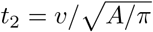) in order for the local order to be propagated beyond immediate neighbours. This finding underlines an important difference in alignment dynamics between active, self-propelling matter, and passive, non self-propelling matter: Self-propulsion will generally increase the number of different neighbours a given cell will interact with, while without self-propulsion there will be fewer interaction partners, but potentially longer interaction times.

**Figure 3:**
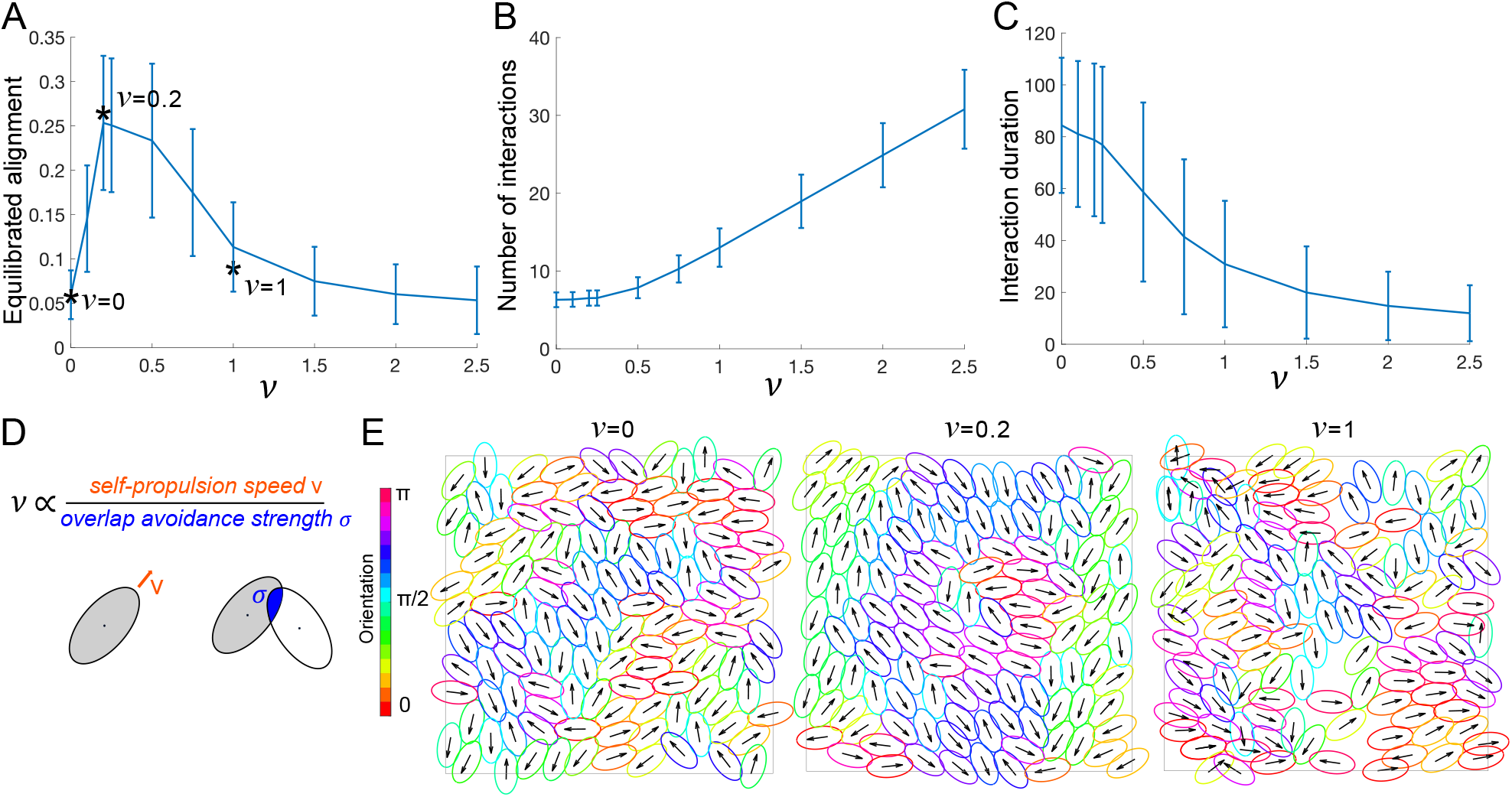
A: Alignment parameter measured at the dynamic equilibrium as a function of *ν*, averaged over 80 simulations, error bars show standard deviation, stars mark simulations in E. B,C: Number of interaction partners per cell (B) and duration of each pair-wise cell interaction (C) between *t* = 0 and *t* = 10 plotted against *ν*, averaged over all 125 cells and 5 simulations. Error bars show standard deviation. D: Schematic explanation of *ν*. E: Simulation snapshots at final time for *ν*-values marked with a star in A. Colors and arrows as in Fig. 2. Fixed parameters: *r* = 2, *N* = 125, *L* = 20.

## 3 Modelling shape changes

### 3.1 Biological motivation

Cells vary in their ability to change shape. Many bacteria, for example, are surrounded by a stiff cell wall, leading to few shape deformations. Most other cell types, including fibroblasts, are soft and deformable, and change their shape dynamically due to internal changes, or in reaction to their surroundings. For example, they can get squished together when confined or become elongated when attached. While cells can also change shape in the absence of other cells, here we only consider shape changes in reaction to interactions with other cells.

### 3.2 Model derivation

We investigate the effect of allowing cells to dynamically change their shape in response to overlap. A summary of model parameter names and meaning can be found in S1 Appendix, Tab. 1. We restrict allowed cell shapes to changing the aspect ratio *r* = *a/b*, where *r*(*t*) is now a function of *t*, while maintaining a constant cell area *A*. To avoid unrealistically large (or small) aspect ratios, we add a term to the energy that punishes deviations from some preferred aspect ratio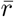, which we set to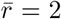. In other words, in the absence of interactions, cell shape will relax towards having the preferred aspect ratio. The strength of this relaxation is given by *g*. This adds an extra term *E*_shape_ to the *E*_tot_ of the base model given in (2),

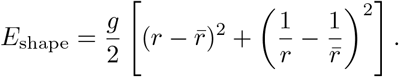

Note that *E*_shape_ is symmetric with respect to 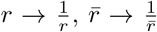. This is to ensure that the shape relaxation behaviour is the same along both axes. The model derivation now follows the same steps as before, for some more details see S1 Appendix, Sec. 1. The equations for **X** and *α* remain unchanged by this. For one cell, we obtain the following equation for how the aspect ratio *r* changes over time:

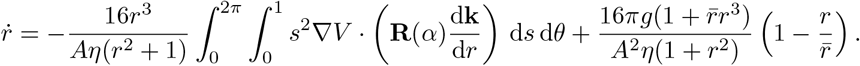

Using the same overlap potential as defined above, we obtain (written in non-dimensional form)

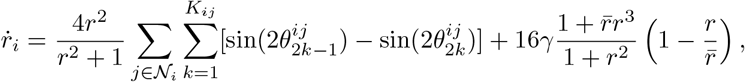

where 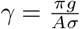 and 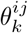 corresponds to the *θ* value that parameterises the overlap point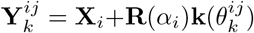.

#### Interpretation for two cells

In the case of the interaction between only two cells with two overlap points (as in Fig. 4C) we have

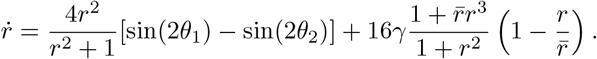

**Figure 4:**
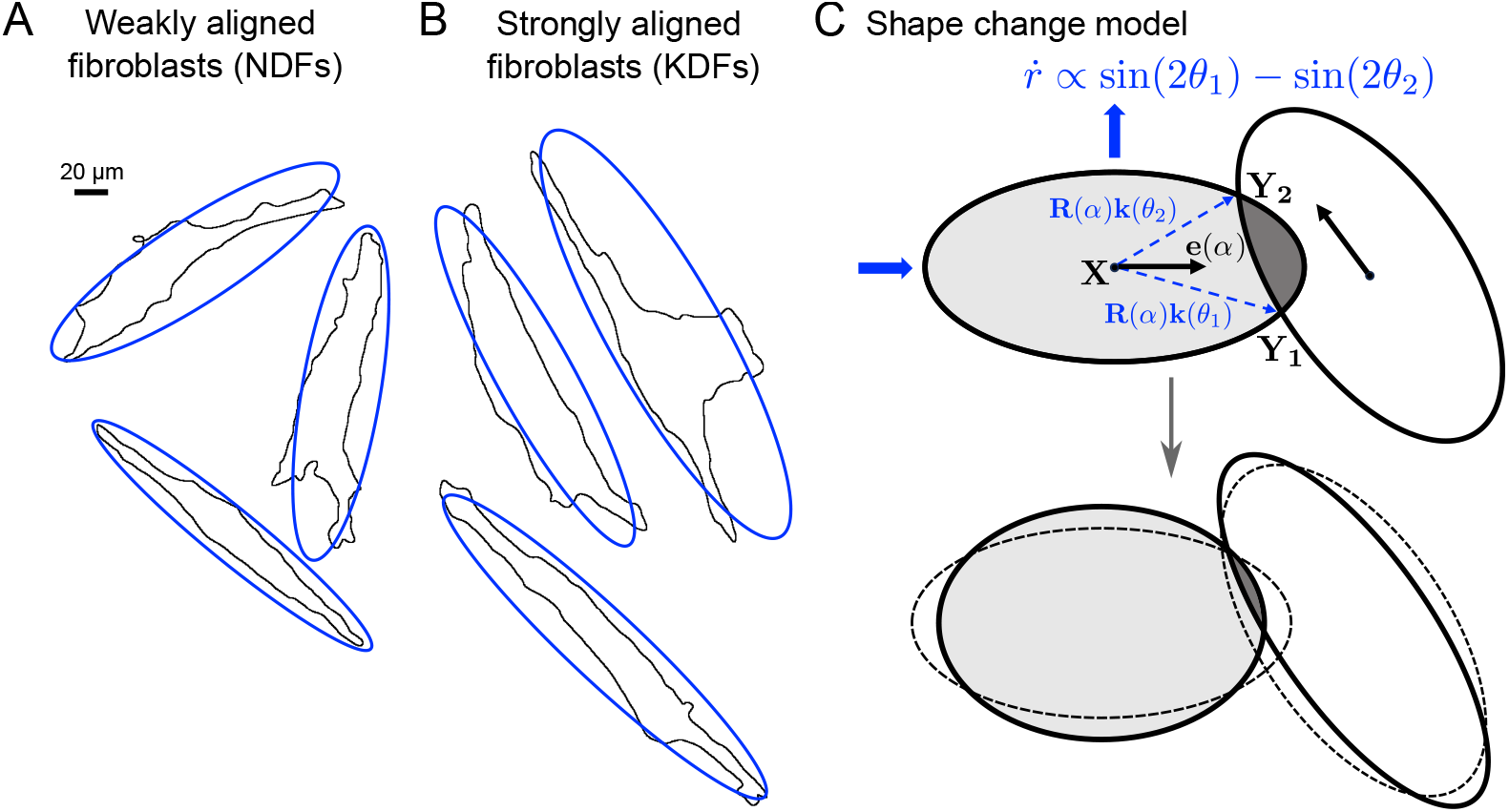
A,B: Single NDF (A) and KDF (B) cells show a range of different shapes in confluent cultures. No difference can be observed between the two samples [47]. C: Schematic of effect of shape change model.

If, as in Fig. 4C, we have that 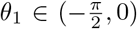 and 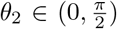, then sin(2*θ*_1_) *−* sin(2*θ*_2_) *<* 0 and the first term will cause the aspect ratio *r* to decrease. This is a result of the cell shortening to avoid overlap. The second term will always act to restore the aspect ratio towards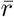. Note that there is only one additional parameter, *γ*, quantifying the restoring force, but no extra parameter quantifying the initial deformation. The reason for this is that shape changes are also driven by minimising overlap and hence based on the same energy term, *E*_overlap_, as the other dynamics (turning and non-propulsion driven translations) driven by overlap avoidance.

### 3.3 Results 2: The base model with shape changes

#### Deformable cells lead to increased alignment

We investigate the effect of *γ*, the strength of the shape restoring force. Small *γ* means cells are more easily deformable, while the limit *γ → ∞* corresponds to non-deformable cells with fixed aspect ratio 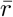 (corresponding to the base model discussed in Sec. 2). Fig. 5A shows that higher deformability correlates with more alignment, suggesting that shape changes aid alignment. Further, we found that allowing cells to deform more increases the average aspect ratio in the population, Fig. 5B. We also assessed whether the optimal value of *ν* found in Fig. 3A is affected by deformability. Fig. 5C shows that indeed, maximal alignment is now reached for larger values of *ν*, i.e. bigger self-propulsion speeds or smaller overlap avoidance.

**Figure 5:**
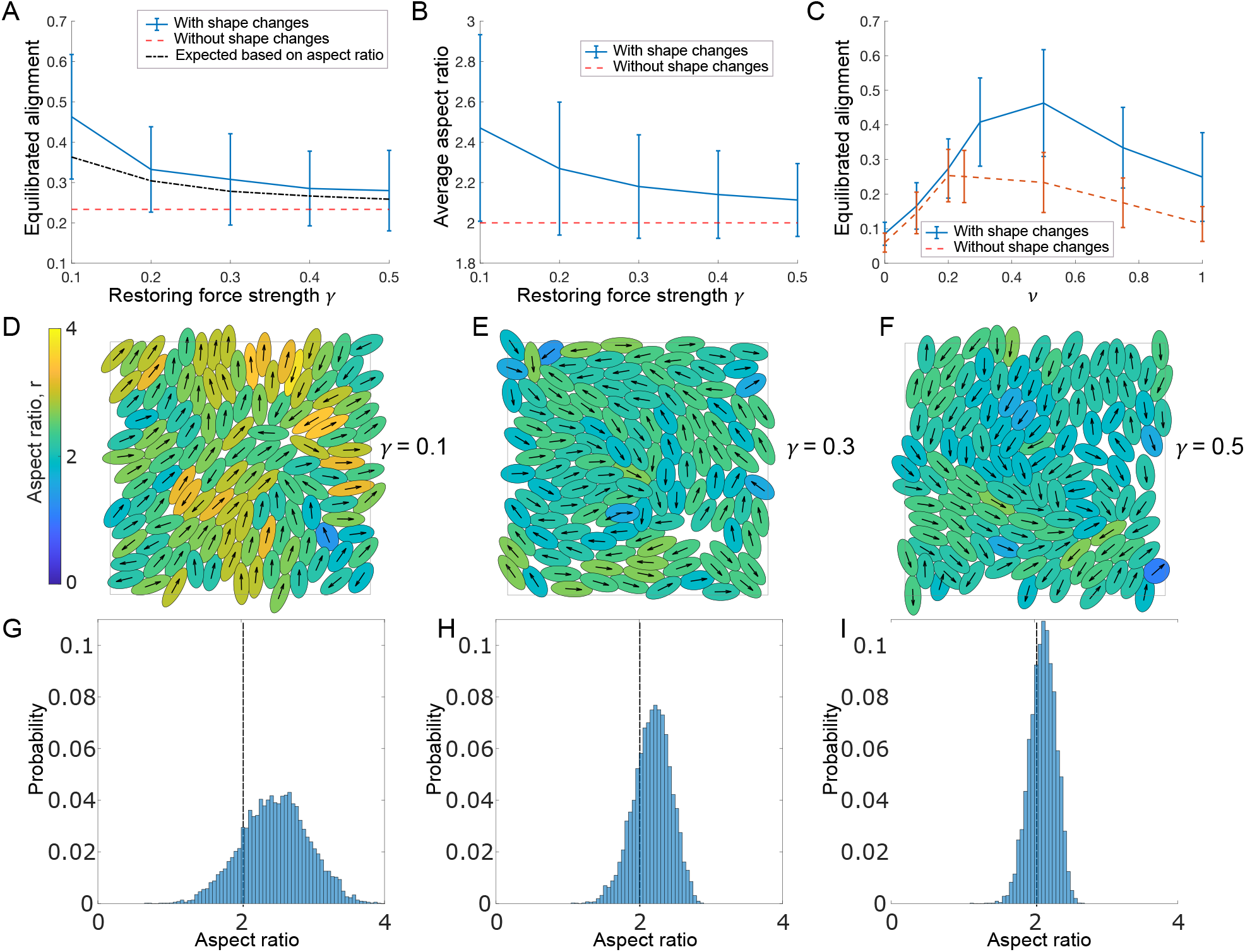
A,B: Equilibrated alignment (A) and average aspect ratio (B) measured for different values of *γ* and compared to non-deformable cells with *ν* = 0.5. In A: Black, dash-dotted line shows expected alignment based on the measured average aspect ratio (see text). C: Equilibrated alignment for different values of *ν* for deformable and non-deformable cells with *γ* = 0.1. D-F: Simulation snapshots at equilibrium for three different values of *γ*. See also S3 Video. Colors indicate aspect ratio. G-I: Distribution of aspect ratio measured for *γ*-values in D-F using values pooled from 60 simulations at equilibrium. Fixed parameters: *N* = 125, *L* = 20,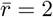.

#### Increased aspect ratio only partly explains increased alignment

We speculated that the increase in alignment for deformable cells is driven by the increase in average aspect ratio. To gain more insight, we inspected the full distribution of aspect ratios for several values of *γ* in equilibrated populations, Fig. 5G-I. We found that while some cells become more rounded (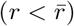), deformability mostly leads to elongated cells (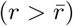). Given that we have already established that higher aspect ratios lead to more alignment (Fig. 2C,D), we tested whether the increase in alignment can be fully explained by the increase in mean aspect ratio. We therefore compared the measured alignment values for two sets of simulations: 1. Deformable cells, for which we determined the mean aspect ratio of the population, Fig. 5A, blue, solid line, and 2. Non-deformable cells with an aspect ratio fixed to the mean aspect ratio of the first group, Fig. 5A, black, dash-dotted line. Interestingly, allowing for dynamic shape changes increases the alignment beyond what the mean aspect ratio would indicate. This suggests that allowing cells to deform dynamically as a reaction to overlap introduces a more flexible way to use the available space leading to higher alignment.

## 4 Modelling cell-cell junctions and actin forces

### 4.1 Biological motivation

In some types of cells, neighbouring cells can form cell-cell junctions. These provide a mechanical coupling as well as a way for cells to exchange signals, i.e. communicate. We model two potential effects of cell-cell junctions: 1. Elastic, reversible connections between two points on the edges of neighbouring cells. 2. Formation of supracellular actin bundles that lead to a bending force. The actin cytoskeleton is the main force generator for moving cells and plays a major role in determining cell shape and polarity. In [47] we observed that the actin network within a cell shares the orientation of the cell itself, with the major bundles typically running along the long axis. Further we found that for keloid fibroblasts (KDFs), neighbouring cells can form supracellular actin bundles, i.e. that the actin bundles of neighbouring cells visually appear to be connected to each other in a smooth manner, without an abrupt change in direction of the actin bundles at the cell-cell junction. This suggests a potential mechanical linkage mediated by cell-cell junctions that causes cells to align their cytoskeleton, and therefore themselves. We will focus on the latter, including elastic connections as a means for these supracellular actin bundles to form. Since the actin bundles are predominantly found to run along the length of each cell, we will only consider cell-cell junctions forming at the front and back of each cell.

### 4.2 Model derivation

A summary of model parameter names and meaning can be found in S1 Appendix, Tab. 1.

#### Elastic connections

In principle cell-cell junctions could form whenever two points in the domain of the ellipses are in close contact. However, since we are interested in cell-cell junctions as a way to mediate the formation of supracellular actin bundles running parallel to the long axis of the cell, we will focus on cell-cell junctions at the front and the rear of cells. We therefore restrict ourselves to only allowing front-to-back, front-to-front and back-to-back junctions. We model these connections as Hookean springs with rest length zero. In principle such connections could be created and broken stochastically, with a distance (or force) dependent breakage rate. However these processes likely happen on a much faster timescale than the overall alignment dynamics. We therefore assume deterministic springs that form and stay in place whenever the distance between connection points are below some critical distance, and break when stretched beyond that distance.

#### Trans-cellular actin cables

Motivated by the biological findings in [47], summarised above, we model the actin network within two connected cells as one rod with a given bending stiffness. For a general rod discretised with a uniform step length *q*, resulting in the points **x**_*i*_ *∈* ℝ^2^, *i* = 1, …, *K*, the bending energy of strength *m* is given by

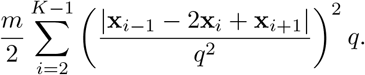

Note that its continuous counterpart would have an integral instead of the sum and the norm of the second derivative instead of the quotient. We discretise the supracellular actin bundle using three points, the two end points not involved in the cell junction, plus the midpoint of the cell-cell junction points. Further we use *q ≈* 2*a*. This formulation has the advantage that it leads to a very simple bending energy. For example, if the front of a cell positioned at **X**_1_ with orientation *α*_1_ is connected to the rear of a cell positioned at **X**_2_ with orientation *α*_2_, the corresponding bending energy would be

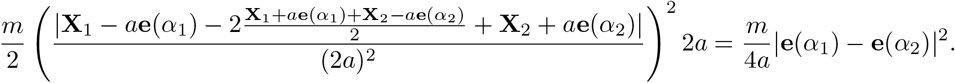

This shows that the bending will only affect the cells’ orientation, not their positions.

#### Incorporation into the full model

We denote the strength of the Hookean springs describing the cell-cell junctions by *k* and assume junctions will exist whenever potential connection points are within distance *l* of each other. To distinguish between front and back connections we define the front and back ends of a cell by **X**^*±*^ := **X** *± a***e**(*α*), and the two relevant index sets by

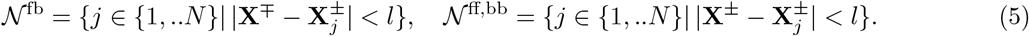

The set *𝒩* ^fb^ describes front-to-back junctions and *𝒩* ^ff,bb^ describes front-to-front and back-to-back junctions. The new contribution to the total energy for a cell positioned at **X** with orientation *α* now takes the form

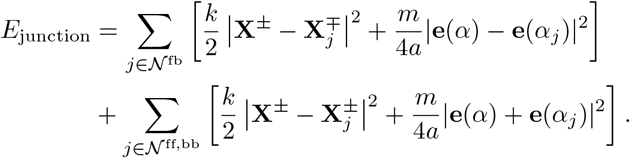

The model derivation now follows the same steps as described in Sec. 2 and yields, in non-dimensional form,

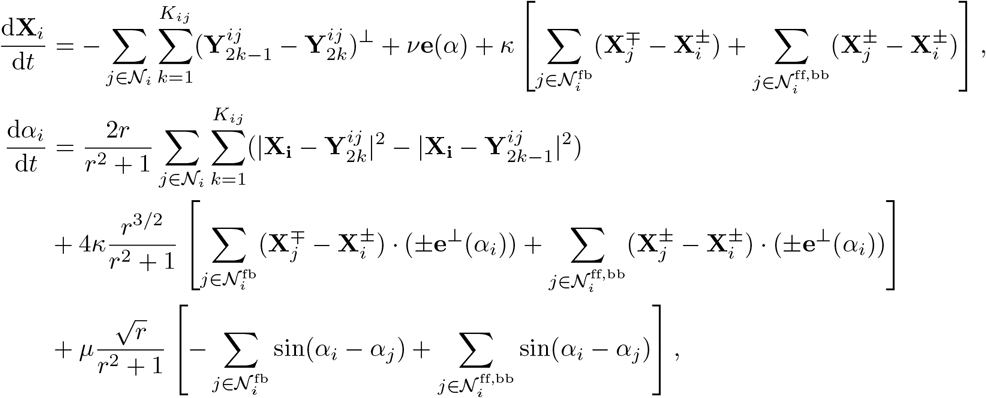

where the index sets *𝒩* ^fb^ and *𝒩* ^ff,bb^ are as defined in (5) with *l* replaced by the non-dimensional 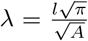 Further we have defined the two non-dimensional quantities 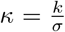 and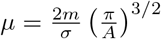, which compare the junction strength and bending strength to the strength of overlap avoidance. We assume that cell aspect ratio, *r*, is constant.

#### Interpretation for two cells

To understand the effect of the new terms, we can return to the situation from above, where the front of cell 1 has a junction with the back of cell 2 (see Fig. 6C,D). Dropping all other terms, this leads to

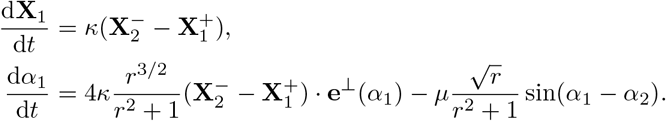

**Figure 6:**
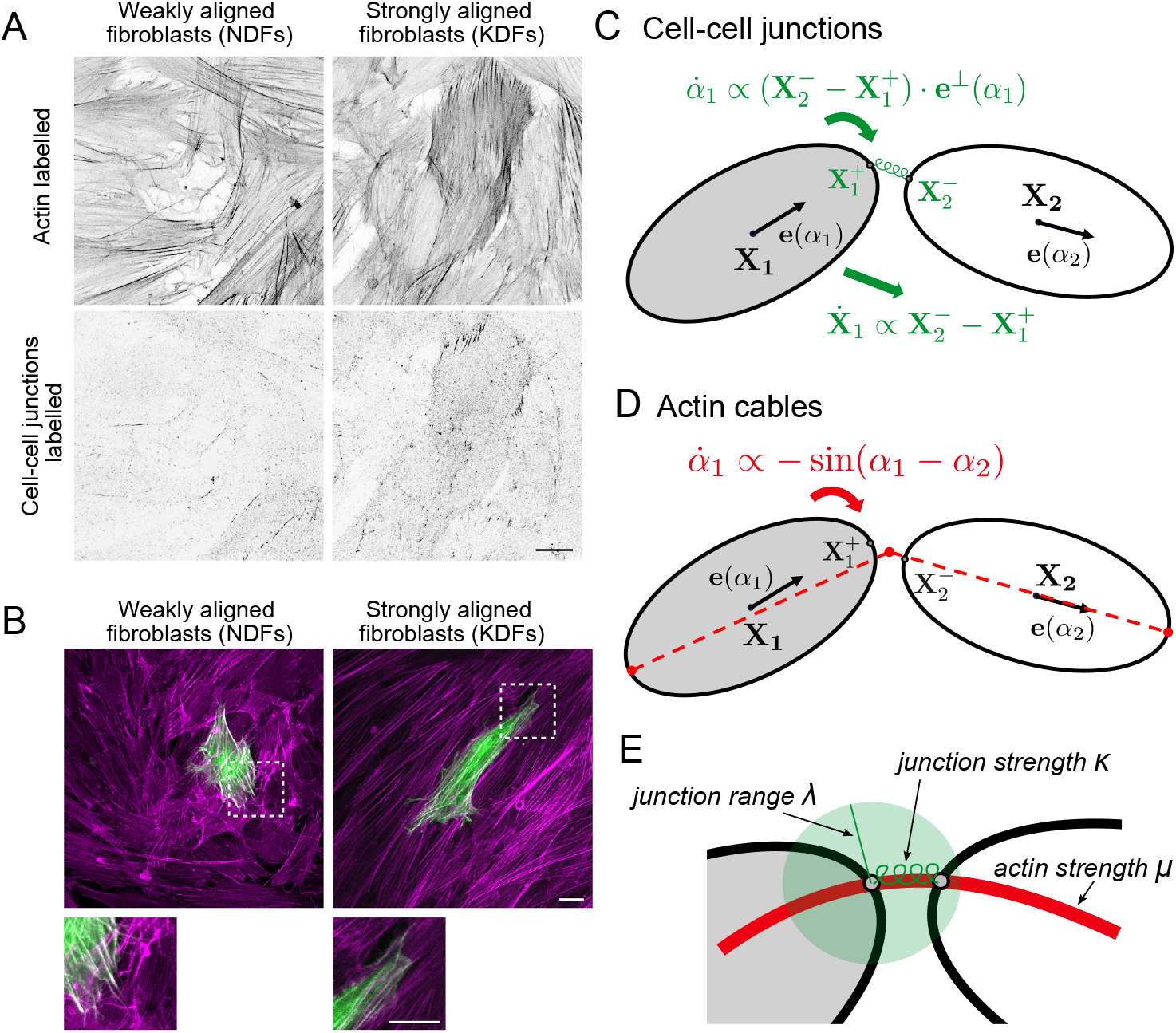
A: NDFs and KDFs stained for F-actin and N-cadherin (a marker for a type of cell-cell junction), revealing an enhanced localisation of N-cadherin at cell-cell junctions in KDF. Scale bar 20 *μm*. B: Supracellular actin bundles. NDF and KDF cultured in vitro for 48 hours, mosaically transfected with EGFP-LifeAct (green) and stained for F-actin (magenta). KDF display more aligned actin bundles spanning multiple cells. Scale bar 20 *μm*. C,D: Schematics showing how the cell-cell junctions (C) and actin forces (D) affect changes in cell position and orientation. E: Explanation of new non-dimensional parameters *λ, κ* and *μ*.

Inspecting the sine-term in the equation for *α*_1_, we see that the cytoskeletal coupling always aids alignment, however larger aspect ratios lead to slower alignment. For the effect of the cell-cell junctions, we see that they cause the center of cell 1 to be pulled along the vector connecting the two junction points. Further, the junction also causes cell turning, however it is not obvious whether this will aid or hinder alignment. The answer becomes even less clear in a multi-cell context and in the context of overlap avoidance and self-propulsion. For this we turn to simulations.

### 4.3 Results 3: The base model with cell-cell junctions and actin forces

#### Cell-cell junctions alone hinder alignment

For cell-cell junctions forming at cell heads and tails there are two new (non-dimensional) parameters introduced to the model: The junction range *λ* (i.e. maximal length over which junctions can form) and the junction strength *κ*. We varied *λ* between 0 and 0.8 (i.e. between 0-28% of one cell length) and *κ* between 0 and 2.5 (i.e. between 0-2.5 times the strength of overlap avoidance), initially without actin forces. First we measured the number of junctions formed and found, as expected, that more junctions are formed as *λ* or *κ* increase, Fig. 7A,C. Next, we found that alignment decreases in both cases, Fig. 7B,D. It appears that the junctions hinder alignment, because they lead to cells forming clumps where more than two cells are joined at one point, which acts against alignment. Indeed, we found many more instances of cell clumps for larger junction range than for lower junction range, as demonstrated in Fig. 7H.

**Figure 7:**
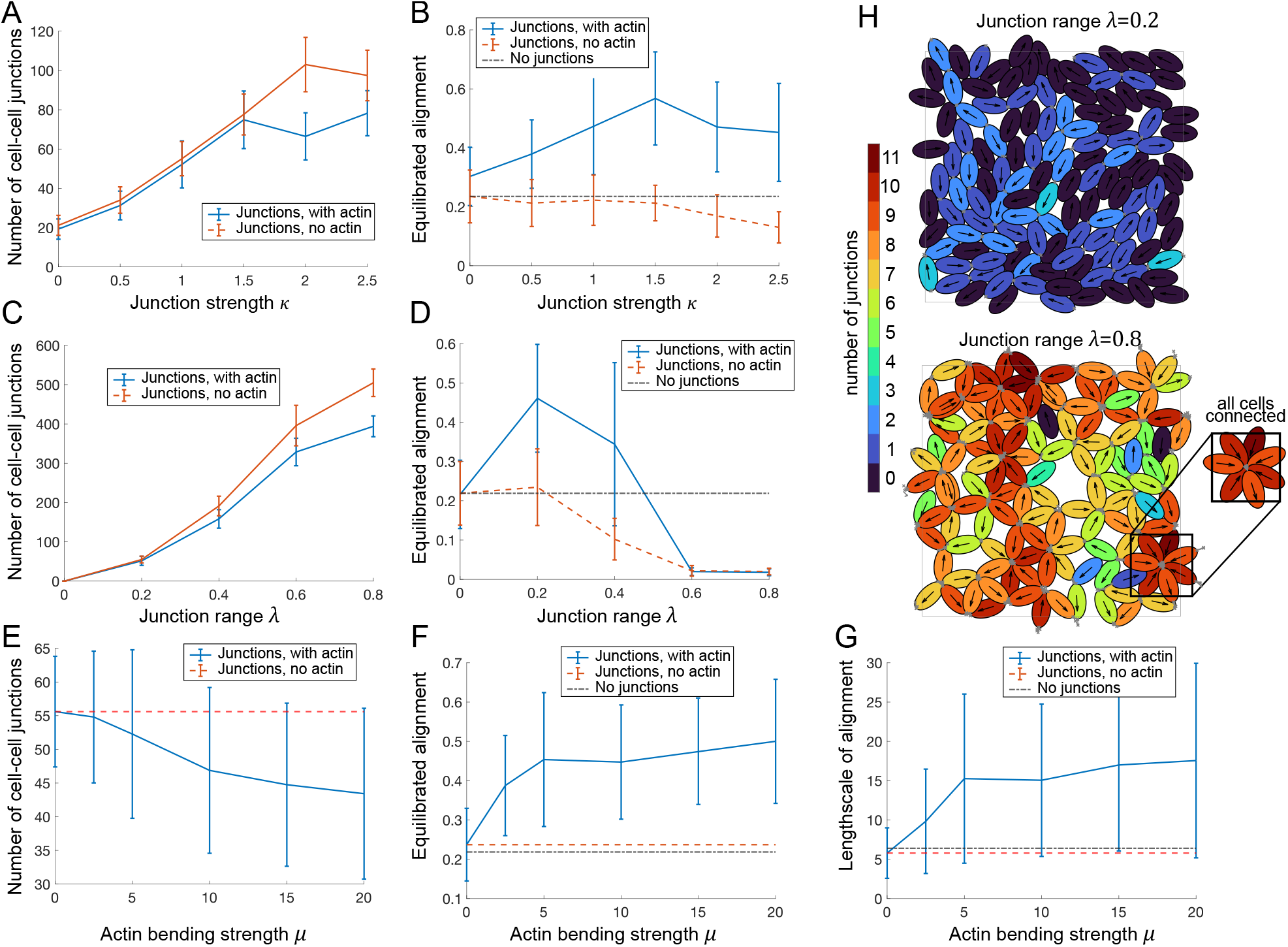
A-F: Number of cell-cell junctions (A,C,E) and value of equilibrated alignment (B,D,F) in dependence of the junction strength *κ* (A,B), the junction range *λ* (C,D) and the actin force *μ* (E,F) with and without an actin force (*μ* = 0 and *μ* = 5 respectively). Average over 60 simulations, error bars show standard deviation. G: Length scale of alignment in dependence on actin force *μ* (see S1 Appendix, Sec. 2). Base parameters are *λ* = 0.2, *ν* = 0.5, *κ* = 1, *N* = 125, *L* = 20. H: Simulation snapshots at equilibrium for *λ* = 0.2 (top) and *λ* = 0.8 (bottom). Cell junctions are marked in grey, color corresponds to number of junctions per cell. Other parameters are *ν* = 0.5, *μ* = 0, *κ* = 1, *N* = 125, *L* = 20.

#### Supracellular actin can greatly aid alignment

Next we investigated how adding an actin force of strength *μ*, representing the effect of supracellular actin bundles, would affect the dynamics. We found that the actin force has only a small effect on the number of junctions formed, with a slight tendency to reduce the number of junctions formed, Fig. 7A,C,E. However, we found that the actin force can greatly increase the equilibrated alignment, Fig. 7B,D,F. We found that as actin forces increase, so does alignment, with alignment values plateauing for large actin forces, Fig. 7F. Further we found that in the presence of actin, there is an optimal junction strength (at *κ ≈* 1.5, i.e the junction strength is 1.5*×*the strength of overlap avoidance), Fig. 7B, and an optimal junction range (at *λ ≈* 0.2, corresponding to about 7% of one cell length), Fig. 7D. Further we found that not only the value of the alignment parameter, but also its length scale, measured as defined in S1 Appendix, Sec. 2, Eq. (2), increases from around 5 (corresponding to 1-2 cell lengths) up to about 17 (corresponding to about 6 cell lengths). In Fig. 7F,G we see a large variation in the alignment values and length scales measured for a given parameter set. We speculated that the reason might be differences in the populations’ junction structure. This we explored next.

#### High alignment at long length scales is driven by linear chains of cells

To understand the junction structure of a cell population better, we represented the simulated cell populations as graphs, where cells and connections between them are represented by the nodes and edges of the graph respectively. This allows for a visual representation of the population structure as well quantification of graph properties, such as the number of junctions per cell (the degree of the node). For a cell to be of degree 0 means it has no connections to other cells, a degree 1 cell has a connection to one other cell, etc. For a given parameter set, we then produced 60 repetitions of the same numerical experiments (with random initial conditions) and inspected, at equilibrium, the correlation between the % of cells of a given degree with the alignment, Fig. 8A-C. Strikingly, we found that for alignment, there is a strong negative correlation with % of degree 0 cells, a strong positive correlation with the % of degree 2 cells, and almost no correlation with the % of degree 1 cells. We saw exactly the same correlation trends for the alignment length scale (not shown). This means for high alignment and high length scales of alignment one needs few unconnected cells and many cells being connected to exactly 2 other cells. In example simulations, Fig. 8D,E we see that, indeed, degree 2 cells form long chains that explain both the high alignment and the long alignment length scales.

**Figure 8:**
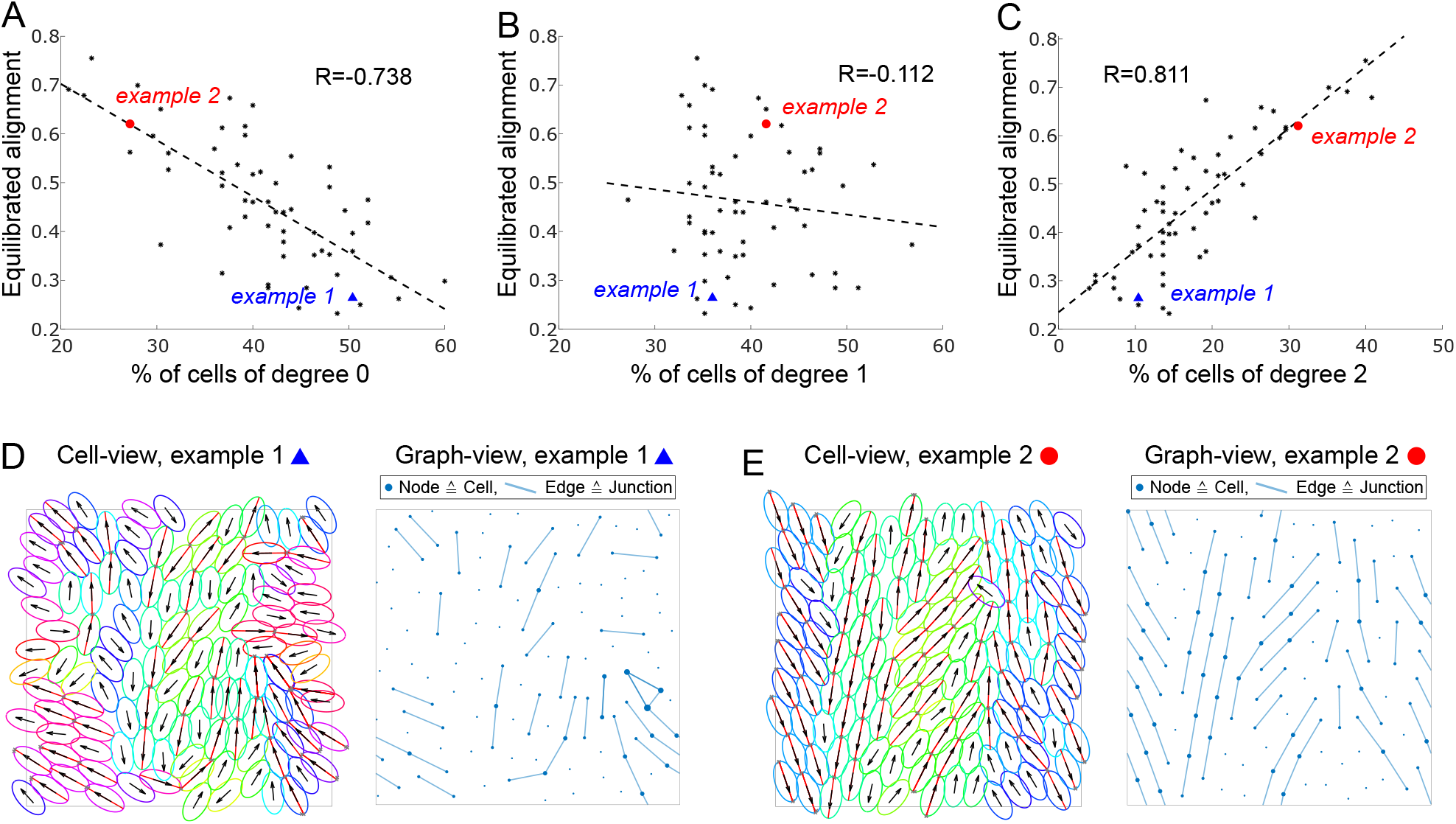
A-C: Scatter plot of equilibrated alignment against % of cells of degree 0 (A), degree 1 (B) and degree 2 (C) at the final time point *T* = 400. Each black dots represents one simulation run. The blue triangle and the red dot mark the examples in D and E respectively. R gives the correlation coefficient and the dotted line gives the linear least squares fit. D,E: Examples marked in A-C in cell-view (left, red lines mark actin, colors and arrows as in Fig. 2) and graph-view (right, dots mark cells/nodes, lines mark edges/junctions). See also S4 Video. Other parameters are *ν* = 0.5, *μ* = 5, *κ* = 1, *λ* = 0.2, *N* = 125, *L* = 20.

## 5 Discussion

### A flexible modelling framework

In this work we have developed a framework to mechanistically model the collective behaviour of active, elliptically shaped particles, that self-propel, avoid overlap, deform and form cell-cell junctions that communicate cytoskeletal forces. The framework is based on energy minimisation and can easily be extended or adapted to include e.g. other cell shapes or different types of cell-cell interactions. A strength of the framework is that the derived equations strike a useful balance between being complex enough to capture the desired phenomena, while being simple enough to be interpretable.

We simulated the emerging collective dynamics for a large ensemble of cells and analysed the numerical results with an emphasis on alignment dynamics. The main computational findings of this work are:

- **Cell alignment needs a balance of self-propulsion and overlap avoidance**. We found that, to maximise collective cell alignment, there is an optimal ratio of self-propulsion speed and overlap avoidance. This allows cells to have a sufficiently long contacts with a sufficient number of cells, which aids alignment.
- **Deformability aids collective alignment**. We found that allowing for flexible cell shapes can aid alignment. We hypothesise that this is because it leads to to more elongated cells (which are associated with more alignment) and a more flexibility of the use of space.
- **Cell-cell junctions alone hinder alignment**. We found that modelling spring-like cell-cell junctions at the cell heads and tails hinders alignment. The reason seems to be the formation of clumps of cells.
- **Actin forces lead to strong, long-scale alignment**. If actin forces are communicated via the cell-cell junctions, this can significantly increase alignment. In this case alignment will happen on a much larger length scale. The reason seem to be long, linear chains of connected cells

### Future work

The derived equations can be used to study e.g. the interaction of only two cells in more depth: Such a simplified system could then be analysed using analytical methods, such as stability analysis, asymptotics or determining long-term behaviour. This is the subject of current ongoing work. The results will give further insights into the involved time scales of movement, the role of self-propulsion or the effect of deformability. In terms of modelling, we are planning on following several directions, such as: 1) We will extend the cell-cell junction model to investigate the effect of cell-cell junctions forming along the whole cell surface. 2) We will investigate how cell-cell junctions affect cell shape. 3) We will derive and analyse a more detailed model of cytoskeletal dynamics within the cell and its interaction with the substrate. These are just some examples. Further, we will test our insights in an experimental setting: Our work in [47], where we compared two types of fibroblasts with different overlap avoidance, is a first step in this direction. However, we will also experimentally test several of the other theoretical predictions in this work.

## 6 Material and methods

For more detailed biological methods, please refer to [47]. The collection of normal skin and keloid scar tissue from patients providing informed consent was ethically approved by the National Research Ethics Service (UK) (14/NS/1073). The study was conducted in accordance with the ethical standards as set out in the WMA Declaration of Helsinki and the Department of Health and Human Services Belmont Report. Normal dermal fibroblasts and keloid derived fibroblasts were isolated from the collected tissue and cultured in vitro. To visualise cell overlap, cells were labelled with CellTrace reagents in two colours (Violet and CFSE, ThermoFisher). Subsequently, cells were plated on imaging substrates with a ratio of 9:1 Violet:CFSE and fixed after 24 hours. Imaging was obtained using a Zeiss LSM 880 confocal microscope (20x NA 0.8 Plan-Apochromat air objective). To evaluate cell shape, single fibroblasts were observed within confluent monolayer cultures. Mosaic expression to highlight single cells was obtained by transfecting cells with EGFP-LifeAct, followed by fixation after 48 hours, and immunostaining with phalloidin to visualise F-Actin (Life Technologies). To visualise cell-cell adhesions, cells were labelled with N-cadherin (Cell Signaling Technologies) after fixation. Imaging was obtained using a Zeiss LSM 880 confocal microscope (40x NA 1.3 Plan-Apochromat oil objective, 40x NA 1.1 LD C-Apochromat water objective, or 63x NA 1.4 Plan-Apochromat oil objective).

## Data Availability Statement

All code written in support of this publication is publicly available at https://github.com/angelikamanhart/Code_Alignment_Ellipses

## 7 Supporting information

**S1 Appendix:** Derivation and computational details.

**S2 Video:** Video corresponding to snapshots shown in Fig. 2A. Parameters are *ν* = 0.2, *N* = 125, *L* = 20, *r* = 2, *T* = 800, time step = 0.01.

**S3 Video:** Video corresponding to snapshot shown in Fig. 5D. Parameters are *ν* = 0.5, *γ* = 0.1, *N* = 125, *L* = 20, 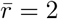, *T* = 400, time step = 0.01.

**S4 Video:** Video corresponding to snapshot shown in Fig. 8E. Parameters are *ν* = 0.5, *λ* = 0.2, *μ* = 5, *κ* = 1, *N* = 125, *L* = 20, 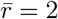, *T* = 400, time step = 0.01.

## 8 Funding and acknowledgements

This work was supported by the Engineering and Physical Sciences Research Council (grant numbers EP/N509577/1, EP/T517793/1), the Wellcome Trust (grant number 107859/Z/15/Z), the European Research Council (ERC) under the European Union’s Horizon 2020 research and innovation program (grant agreement no. 681808) and BBSRC project grant (BB/V006169/1). The authors wish to thank Antoine Nicolas Diez for his helpful thoughts and comments.

