## Supplementary material for "Derivation and simulation of a computational model of active cell populations: How overlap avoidance, deformability, cell-cell junctions and cytoskeletal forces affect alignment": S1 Appendix

#### 1 Model derivation details

**The base model.** Here we detail the model derivation steps for $\frac{\mathrm{d}\mathbf{X}}{\mathrm{d}t}$. The other equations can be derived analogously. The energy for one cell is given by

$$E_{\Delta t} = \int_0^{2\pi}\int_0^1 \left[\eta\frac{|\mathbf{x}(t,s,\theta)-\mathbf{x}(t-\Delta t,s,\theta)|^2}{2\Delta t} - \mathbf{F}\cdot\mathbf{x}(t,s,\theta) + V(\mathbf{x}(t,s,\theta))\right] abs\,\mathrm{d}s\,\mathrm{d}\theta.$$

Step 1: Taking the derivative with respect to $\mathbf{X}$. We hold $r(t-\Delta t,s,\theta)$ constant and obtain

$$\frac{\mathrm{d}E}{\mathrm{d}\mathbf{X}} = \int_0^{2\pi}\int_0^1 \left[\eta\frac{(\mathbf{x}(t,s,\theta)-\mathbf{x}(t-\Delta t,s,\theta))}{\Delta t} - \mathbf{F} + \nabla V(\mathbf{x}(t,s,\theta))\right] abs\,\mathrm{d}s\,\mathrm{d}\theta.$$

Step 2: Setting $\frac{\mathrm{d}E}{\mathrm{d}\mathbf{X}} = 0$ letting $\Delta t \to 0$. Using Eq. (1) in the main text to determine $\dot{\mathbf{X}}$ we obtain

$$0 = \int_0^{2\pi}\int_0^1 \left[\eta(\dot{\mathbf{X}} + s\dot{\alpha}\frac{\mathrm{d}\mathbf{R}}{\mathrm{d}\alpha}\mathbf{k}) - \mathbf{F} + \nabla V(\mathbf{x}(t,s,\theta))\right] abs\,\mathrm{d}s\,\mathrm{d}\theta.$$

Step 3: Evaluation of the integrals. The first term gives $A\eta\dot{\mathbf{X}}$, corresponding to the (equal) friction experienced across the cell area. The second term integrates to zero, due the cosine and sine terms in $\mathbf{k}$. The third term involving self-propulsion simply gives $A\mathbf{F}$, which we set to be $\mathbf{F} = v\eta\mathbf{e}(\alpha)$. Together this yields the first equation in (3) in the main text. For the overlap avoidance term, we start by reverting to Cartesian coordinates and using the divergence theorem.

$$\int_0^{2\pi}\int_0^1 abs\nabla V(\mathbf{x}(t,s,\theta))\,\mathrm{d}s\,\mathrm{d}\theta = \int_{\mathcal{A}} \nabla V(\mathbf{x})\,\mathrm{d}\mathbf{x} = \int_{\partial\mathcal{A}} V(\mathbf{x})\mathbf{n}\,\mathrm{d}S,$$

where $\mathbf{n}$ is the outward unit normal along the edge of the cell. Next we assume there is only overlap with one other cell covering the domain $\mathcal{B}$ and denote the intersection points going counter-clockwise along the intersecting segment by $\mathbf{Y}_1$ and $\mathbf{Y}_2$. We use the piecewise constant choice of $V = \mathbb{1}_{\mathcal{A}\cap\mathcal{B}}$. This gives

$$\int_{\partial\mathcal{A}} V(\mathbf{x})\mathbf{n}\,\mathrm{d}S = \sigma\int_{\partial\mathcal{A}\cap\mathcal{B}} \mathbf{n}\,\mathrm{d}S = \sigma\left(\mathbf{Y}_1 - \mathbf{Y}_2\right)^{\perp}.$$

Note that the fact that we can evaluate the overlap avoidance integral explicitly hinges on the simple shape of $V$.

**Modelling shape changes.** We follow the derivation of the base model, noting that there is an additional term, $E_{\text{shape}}$, in the energy. We then take the derivative with respect to $r$ and note that $a$ and $b$ are no longer constant, but that the area of the cell $A = \pi ab$ does remain constant. The equations for $\dot{\mathbf{X}}$ and $\dot{\alpha}$ remain unchanged. For $\dot{r}$ we obtain

$$0 = \frac{A^2\eta(1+r^2)}{16\pi r^3}\dot{r} + \frac{A}{\pi}\int_0^{2\pi}\int_0^1 s^2\nabla V \cdot \left(\mathbf{R}(\alpha)\frac{\mathrm{d}\mathbf{k}}{\mathrm{d}r}\right)\,\mathrm{d}s\,\mathrm{d}\theta + g(r-\bar{r})\frac{1+\bar{r}r^3}{\bar{r}r^3}.$$

On evaluating the integral, again using the divergence theorem, and using that $\frac{\mathrm{d}\mathbf{k}(\theta)}{\mathrm{d}r} = \frac{1}{2r}\mathbf{k}(-\theta)$, we obtain the governing equation for $\dot{r}$.

### 2 Computational details

| Biological parameters | | |
|---|---|---|
| **Parameter** | **Dimensions** | **Comment** |
| $N$ | | Number of cells in the domain |
| $a, b$ | length | Semi major and minor axis of ellipse |
| $A$ | length$^2$ | Area of ellipse (kept constant) |
| $v$ | $\frac{\text{length}}{\text{time}}$ | Self propulsion speed |
| $\eta$ | $\frac{\text{mass}}{\text{length}^2\text{time}}$ | Strength of friction with the substrate |
| $\sigma$ | $\frac{\text{mass}}{\text{time}^2}$ | Strength of overlap avoidance |
| $g$ | $\frac{\text{mass}\times\text{length}^2}{\text{time}^2}$ | Strength of relaxation to preferred aspect ratio |
| $m$ | $\frac{\text{mass}\times\text{length}^3}{\text{time}^2}$ | Strength of bending energy |
| $k$ | $\frac{\text{mass}}{\text{time}^2}$ | Strength of spring/ cell cell junction |
| $l$ | length | Cell-cell junction range |
| Scaled (dimensionless) parameters | | |
| **Parameter** | **Definition** | **Comment** |
| $L$ | $\frac{\text{physical length}}{\sqrt{A/\pi}}$ | Scaled side length of square domain |
| $r, (\bar{r})$ | $a/b$ | (Preferred) aspect ratio of ellipse |
| $\nu$ | $\frac{v\eta}{\sigma}\sqrt{A\pi}$ | Ratio of time scale of overlap avoidance to time scale of self propulsion. |
| $\gamma$ | $\frac{\pi g}{A\sigma}$ | Ratio of time scale of overlap avoidance to time scale of relaxation to preferred aspect ratio |
| $\mu$ | $\frac{2m}{\sigma}\left(\frac{\pi}{A}\right)^{3/2}$ | Ratio of actin force to overlap avoidance strength |
| $\kappa$ | $\frac{k}{\sigma}$ | Ratio of cell-cell junction strength to strength of overlap avoidance |
| $\lambda$ | $l\sqrt{\frac{\pi}{A}}$ | Dimensionless junction range |

Table 1: Table of parameters

**Computational Method & Parameters.** To numerically solve the governing equations we use a Forward Euler method with $\Delta t = 1/100, 1/150$, depending on the parameters. We use $N = 125$ cells and a square domain with $\Omega = [0, L]^2$, $L = 20$ together with periodic boundary conditions. For the initial cell orientations we used a uniform random distribution on $[0, 2\pi)$. Initial cell positions were distributed uniformly on $\Omega$ with a minimum distance of 1 between cells. The only source of stochasticity in the system are the initial conditions. To draw conclusions, we averaged output quantities over 60 or more simulations.

**Finding overlap points.** Numerically solving the governing equations hinges on finding the points of overlap for all ellipses. We use an approximate method to do this, which is computationally more efficient than solving for intersection points analytically and also allows for easier generalisation to other cell shapes. The boundary of each ellipse is discretised by 200 points evenly spaced by angle. For a given ellipse $i$, we then use the `rangesearch` function in MATLAB to find all ellipses within a radius of $2a$ of ellipse $i$. For this set of ellipses that have the potential to be overlapping, we search for all boundary points within a certain optimal search range that is small enough to identify two distinct points of overlap, but large enough to not miss any. This will depend both on the aspect ratio of the ellipse and on the number of points on the boundary. For example, for ellipses of aspect ratio 2, the optimal search range is 0.04. Clusters of neighbouring boundary points are identified and averaged to find an approximate point of overlap and the corresponding $\theta$ value. We determine the ordering of overlap points as described in Tab. 2. Finding the cell cell junction points is simply done by defining the points at the head and tail of each cell and identifying those within the pre-defined junction range $\lambda$ using Matlab's `searchrange` function.

| Number of overlap points | Method |
|---|---|
| 1 | Point is disregarded |
| 2 | For two overlap points $\mathbf{Y}_a$ and $\mathbf{Y}_b$ parameterised by $\theta$ values $\theta_a$ and $\theta_b$, if $(\theta_a - \theta_b) \mod 2\pi < (\theta_b - \theta_a) \mod 2\pi$ then $a = 2$ and $b = 1$, otherwise $a = 1$ and $b = 2$. |
| 3 | We identify the point where the ellipses are most likely to be tangential as the point with the maximum cluster size. This point is disregarded and we proceed as with 2 overlap points. |
| 4 | We order the points in order of increasing $\theta$ as $\mathbf{Y}_a, \mathbf{Y}_b, \mathbf{Y}_c, \mathbf{Y}_d$. We find the vectors between each consecutive pair of overlap points, and then identify the one that is oriented most closely to the orientation $\alpha$, up to a multiple of $\pi$. These two points make up the first pair of overlap points, and the remaining points make up the other. e.g. if $\mathbf{Y}_d - \mathbf{Y}_c$ is the vector closest in orientation to $\alpha$ then $c = 1, d = 2, a = 3, b = 4$. |
| 5 | The two overlap points that are closest together are identified and one is disregarded. We then treat as in the case of 4 overlap points. |

Table 2: Overlap point treatment.

**Alignment Parameter.** Alignment at time $t$ in the neighbourhood $l$ is measured using the formula

$$\text{Alignment}_t(l) = \sqrt{\langle \cos 2\Delta\phi_{l,t} \rangle^2 + \langle \sin 2\Delta\phi_{l,t} \rangle^2}, \qquad (1)$$

where $\Delta\phi_{l,t}$ is the difference in orientation between every pair of cells at time $t$ within the alignment neighbourhood given by a distance of radius $l$. The bracket $\langle\rangle$ denotes the average of all these values for a given group of cells. Note that this is different to how alignment is measured in [47]. All alignment plots in this work show alignment with a neighbourhood size of $l = 6$. To determine the length scale of alignment, we measure $\text{Alignment}_t(l)$ for different neighbourhood sizes $l$ and average over several simulations. We then fit the resulting values to the function

$$\text{Alignment}_t^{\text{fit}}(l) = ae^{-\frac{l}{l_a}}. \tag{2}$$

From this we obtain the length scale of alignment $l_a$, which indicates how quickly in space alignment decays. Larger $l_a$ means slower decay and hence a longer length scale of alignment.

**Packing fraction.** We find the packing fraction by dividing the area of the domain that is covered by cells by the total area of the domain. The higher the packing fraction, the less overlap there is in the population. As a consequence of the chosen reference length, all cells have area $\pi$. Therefore, for $N = 125$, $L = 20$, the maximum packing fraction (which corresponds to no cell overlap) is $\frac{125\pi}{20\times20} \approx 0.98$.

**Cell Interactions.** To quantify cell-cell interactions, we determine the index-pairs of overlapping cells at each time point and follow this set over time. From this, we can calculate 1. The average time of interaction and 2. The average number of interactions per cell at each time point.
